# Conneconomics: The Economics of Dense, Large-Scale, High-Resolution Neural Connectomics

**DOI:** 10.1101/001214

**Authors:** Adam H. Marblestone, Evan R. Daugharthy, Reza Kalhor, Ian D. Peikon, Justus M. Kebschull, Seth L. Shipman, Yuriy Mishchenko, Jehyuk Lee, David A. Dalrymple, Bradley M. Zamft, Konrad P. Kording, Edward S. Boyden, Anthony M. Zador, George M. Church

## Abstract

We analyze the scaling and cost-performance characteristics of current and projected connectomics approaches, with reference to the potential implications of recent advances in diverse contributing fields. Three generalized strategies for dense connectivity mapping at the scale of whole mammalian brains are considered: electron microscopic axon tracing, optical imaging of combinatorial molecular markers at synapses, and bulk DNA sequencing of trans-synaptically exchanged nucleic acid barcode pairs. Due to advances in parallel-beam instrumentation, whole mouse brain electron microscopic image acquisition could cost less than $100 million, with total costs presently limited by image analysis to trace axons through large image stacks. It is difficult to estimate the overall cost-performance of electron microscopic approaches because image analysis costs could fall dramatically with algorithmic improvements or large-scale crowd-sourcing. Optical microscopy at 50–100 nm isotropic resolution could potentially read combinatorially multiplexed molecular information from individual synapses, which could indicate the identifies of the pre-synaptic and post-synaptic cells without relying on axon tracing. An optical approach to whole mouse brain connectomics may therefore be achievable for less than $10 million and could be enabled by emerging technologies to sequence nucleic acids in-situ in fixed tissue via fluorescent microscopy. Strategies relying on bulk DNA sequencing, which would extract the connectome without direct imaging of the tissue, could produce a whole mouse brain connectome for $100k–$1 million or a mouse cortical connectome for $10k–$100k. Anticipated further reductions in the cost of DNA sequencing could lead to a $1000 mouse cortical connectome.

## 1 Introduction

Wiring diagrams for neuronal microcircuits support efforts to reverse-engineer the brain and to identify structural contributors to neuropsychiatric pathologies [1–3]. Acquisition of large-scale connectivity data could, for example, help to guide efforts to simulate emergent network functions in mammalian brains [4], which are currently based on statistical extrapolations from small datasets [5, 6]. Recently, the field of *connectomics* has sought to develop technologies to rapidly extract comprehensive cellular-resolution maps of synaptic connectivity [7].

Multiple toolsets could potentially support connectomics at the scale of entire mammalian brains or brain regions. These include automated electron microscopy and image analysis as well as newer techniques for DNA sequencing of cell-identifying molecular barcode tags [3]. It is unclear, however, to what degree these could be leveraged to create a scalable, integrated connectomics solution, and whether this could be done at a reasonable cost.

Here we analyze the design space for connectomics by considering the scaling and cost constraints on a range of solutions. We focus here on techniques for dense, cellular-resolution circuit mapping of individual brains: we do not consider sparse mapping (e.g., viral tracers), low-resolution mapping (e.g., diffusion MRI) or mapping based on functional measurements [8, 9].

Approaches differ widely in the cost requirement for obtaining the complete connectome of an *individual mammalian brain*, such as the mouse brain, with 7.5 × 10^7^ neurons in a volume of 420 mm^3^ (a large fraction of these are in the cerebellum, roughly 3 × more than in cortex [10]). They also differ in the nature of the additional information which they provide, beyond the abstract cell-cell connectivity matrix.

In Sections 2 and 3, we review the existing electron microscopy approaches, as well as a recently proposed DNA sequencing approach called BOINC [3], focusing on their scalability towards the mapping of large volumes of mouse brain tissue. Finally, in Section 4, we discuss the prospects for connectomics solutions based on direct imaging by optical microscopy.

### 1.1 Challenges for Connectomics

Generating microscale anatomical wiring diagrams is a major technological challenge [11]. To understand why this is the case, we begin by outlining some of the relevant structural features of neural circuits. As discussed in detail below in the context of specific methods, these features place stringent requirements on technologies for comprehensive measurement of synaptic connectivity. Depending on the method used to measure connectivity, different sets of features become critical in constraining the design space.

#### Packing density

Neurons are packed densely in a three-dimensional jungle of wiring: there are roughly 100,000 neurons per mm^3^ and 1–2 synapses per µm^3^ on average inside mouse neocortex. In rat CA1 hippocampal neuropil, the spatial distribution of synapses appears to be consistent with a uniform random distribution on length scales above the synaptic size [12, 13], with a mean synapse-synapse distance of ∼480 nm (see [13] for the measured distribution of distances). Measurements in rat layer III somatosensory cortex also suggested an approximate uniform distribution subject to the constraint that synapses cannot overlap in space [14], again with nearest-neighbor distances of ∼500 nm. If the locations of synapses are distributed uniformly, the number of synapses per cubic micron will conform approximately to a Poisson distribution, with mean density of 1–2 synapses per µm^3^: 13%–37% probability of no synapses, 27%–37% one synapse, 18%–27% two synapses, 6%–18% three synapses, 1.5%–9% four synapses, 0.3%–4% five synapses and 0.05%–1% six synapses.

#### Spatial variability

The spatial density and arrangement of synapses varies by region, cortical layer (see [15] for glutamatergic synapse density vs. layer in mouse neocortex), and so forth, although there appears to be a roughly universal number of neurons *beneath* a square of fixed area, say 1 mm^2^, of the cortical surface, varying by a factor of less than 1.6 in rodents [16]. Furthermore, on some neurons, specific classes of synaptic contacts are spatially organized on the target dendrites [17, 18]. Unfortunately, detailed measurements of these distributions are currently only available for a handful of brain locations.

#### Multiplicity

There is a large variation in the *number* of synaptic contacts between any given connected *pair* of cells. In hippocampus, synaptically connected neurons are often linked by only one synapse, with higher level redundant connectivity occurring in a group of nearby neurons. In some areas of cortex there are only a handful of contacts between synaptically-paired cells [19], while in other areas there can be as many as a dozen or more, e.g., 6 ± 5 (mean ± standard deviation) among thick-tufted neurons in developing rat L5 neocortex [20]. In general these distributions are unknown. At some synapses outside cortex (e.g., the Calyx of held [21]) the effective number of “synapses” (i.e., vesicle release sites) is much higher.

#### Small feature sizes

Relevant anatomical features of neurons are on the nanoscale, below the wavelength of light: dendritic spine necks and axons shrink in diameter down to tens of nanometers. Synapses can be as small as ∼200 nm in diameter (including both pre and post-synaptic compartments) [22].

#### Long projections

Axons often travel several millimeters along complex paths, with *kilometers* of axonal wiring present in a cubic millimeter of cortex. Furthermore, at least a few cubic millimeters of reconstructed volume are likely needed to adequately define the connectivity of local cortical circuits, though smaller volumes may be sufficient to reconstruct canonical circuit patterns in other brain areas [7].

#### Diversity

Mammalian connectomes are not identical across different individuals, so many connectomes should be mapped. Methods for statistical reconstructions of connectomes by combining partial reconstructions from multiple animals [23, 24] can be useful for determining average connectomes as well as statistical variation around the average. To the greatest extent possible, however, multi-modality measurements should be integrated such that they can be simultaneously applied to each individual brain under study, rather than averaging or correlating across different brains. The ideal technique would be sufficiently low cost that many individual connectomes could be rapidly acquired. Post-hoc correlation across multiple single-brain connectomes could reveal insights at the level of mechanistic conservation: for example, there are likely connection motifs which are invariant across individuals, e.g. in the organization of cortical circuits.

#### Size of dataset

The amount of data needed to store the abstract connectivity matrix of a mouse brain is roughly *N* · *s* · log_2_(*N*) = 2.65 × 10^12^ bits *<* 1 terabtye, where *N* ≈ 10^8^ is the number of neurons and *c* ≈ 10^3^ is the average number of synapses per neuron [25]. Including synaptic weights and molecular profiles has been estimated to increase this storage requirement by *<* 100× [26].

#### 1.2 Caveats for Cost Calculations

Below, we attempt to estimate the costs associated with hypothetical whole-mouse-brain connectomics projects – normalized to a three-year project – based on a variety of technology platforms. These estimates are intended as rough approximations and should not be taken literally as proposed figures for particular projects. Despite these caveats, it is of interest to explore how even crude estimates of project cost vary with changes to the technology architecture adopted, or with improvements to particular parameters, such as the speed of super-resolution optical microscopy or the number of parallel electron beams per electron microscope.

## 2 Electron Microscopy (EM) Connectomics

Electron microscopy is the most thoroughly developed approach for the dense reconstruction of neural circuits. Because the wavelength of an electron under 10 kV accelerating voltage is ∼10 pm, imaging with electrons can (in principle) reach spatial resolutions in the sub-nanometer to nanometer range [27], more than sufficient to trace the finest morphological sub-structures of neurons. The basic strategy employed by the current EM approaches is to obtain many morphological images of thin tissue sections, segmenting those images into regions corresponding to distinct neuronal processes, and tracing individual axons from one image to another. Because axons are thin, long, and densely interspersed with other neuronal processes, tracing their entire lengths is a challenge.

### 2.1 EM Data Acquisition: Basic Properties

#### Beam current and bit precision

The physical constraints on large-scale electron microscopy for neural circuit reconstruction were first studied in the 1980s [28], following the acquisition of the *C. elegans* connectome by electron microscopy [29]. The electron dose per pixel is one property which constrains the resolution and speed of an imaging system. An exemplary recent connectomics study used roughly 14 electrons per nm^2^ [30], or 3812 electrons per 16.5 nm × 16.5 nm pixel. Due to Poisson counting statistics, the fractional error in the estimate of the stain density in a voxel goes roughly as 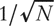, where *N* is the number of electrons passing through the voxel [28], so the analog bit precision in that study was roughly log_2_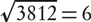 bits at each pixel.

Merkle [28] used the number of electrons per voxel, the number of parallel electron microscopes available, and the total project time to estimate the beam current per microscope: imaging a whole human brain in 3 years at 10 nm × 10 nm × 10 nm × voxel size, with 7-bit precision and 1000 parallel microscopes, would give 0.1 mA beam current, comparable with that of electron microscopes circa 1989.

#### TEM vs. SEM

Transmission electron microscopy (TEM) involves passing electrons *through* a sample, whereas scanning electron microscopy (SEM) relies on back-scattered or secondary electrons emitted from the sample’s surface. High-resolution EM analysis was originally limited to transmission electron microscopy, which necessitated the use of ultra-thin (*<* 100 nm), grid-suspended sections to allow electron penetration through the slice. Although TEM sections cannot easily be made *thinner* than a few tens of nanometers, *z*-resolution can be improved by tilting the sample and performing a tomographic reconstruction [31]; only a handful of additional tilts are required if sparse reconstruction techniques are used [32]. Indeed, the first proposals for whole-mouse-brain electron microscopy circuit tracing [28] assumed a TEM tomography strategy.

#### Scaling up TEM

Unfortunately, large-scale automation of transmission electron microscopy has been difficult in practice due to the need to isolate fragile ultra-thin sections which can be penetrated by the electron beam [33, 34]. TEM is still used today, at rates approaching 10 megapixels per second using camera arrays [35], but in a recent study, ∼30 of ∼4000 thin sections were lost in the preparation process [35]. Thus, improvements in TEM sample handling are needed to trace connectivity at whole-mouse-brain scale, and we focus on scanning electron microscopy techniques below. Improvements in high-throughput, high-reliability automated TEM sample preparation, coupled with camera arrays [35], could make TEM viable for large-scale circuit reconstruction [31].

#### Maximum block size and the importance of lossless subdivision

EM cannot take advantage of parallel imaging on multiple machines unless lossless subdivision of the tissue into “blocks” is performed prior to imaging: it must be possible to separately image two adjacent sub-blocks and stitch the resulting images together in software. The finest neuronal processes must be traceable from one sub-block to the other, and features localized at the block-block interface must be preserved. In one demonstrated technique for lossless subdivision [33, 36], a hot diamond knife reduces the cutting stress locally and reversibly, and an oil film prevents damage due to scraping of the tissue block along the knife edge. This process appears amenable to large-scale automation.

#### Parallel beam instruments

The speed of SEM can be increased by using multiple parallel beams in a single instrument. For example, Zeiss is developing a multi-beam SEM (mSEM) instrument with 61-fold parallelization. It is incorrect to assume, however, that the speed of a multibeam SEM scales proportional to the number of beams. Because of the limitations of electron optics and charge repulsion, the total current in each beam is typically much smaller than can be achieved in a single-beam system. A 10× speed improvement over an equivalent single-beam instrument would be a more conservative estimate, even though the system has 61 beams. Parallelization of a 40 mega-pixel per second SEM by a factor of 25 would lead to gigapixel per second rates, which appears to be a reasonable upper bound for the immediate future. More optimistically, advanced SEMs could potentially use thousands of parallel beams, and instrument costs could be reduced to the $100k regime via solid-state lithographic electron optics [33]; such systems may be a natural offshoot of the development of next-generation electron-beam lithography systems by the semiconductor industry.

#### Reliability and cost of sectioning

Reliability of ultra-thin-sectioning is a key issue for SEM approaches. Empirically, it is currently difficult to knife-section a 300 µm × 300 µm × 300 µm block at 30 nm slice thickness, and usually takes multiple attempts; reliable sectioning becomes more difficult for larger block sizes. We highlight scenarios below where reliability of physical sectioning is likely to become the major limiting factor. Note also that at high electron doses, the mechanical properties of the block surface change in such a way to worsen the minimum section thickness and the sectioning reliability.

Diamond knives used in electron microscopy routinely perform 10k sections before incurring damage. Assuming that only 1000 sections are used per knife to keep damage rates conservatively low, and that each knife costs $2500, the cost of the knives for 420 mm^3^*/*(1 cm^2^ × 25 nm) = 168000 sections would be <$500k.

Another major challenge to whole brain imaging will be minimizing the material loss from vibratome section to vibratome section, and from the sub-sectioning of the brain either before or after embedding.

## 2.2 Approaches to automated SEM

Three strategies for large-scale electron-microscopy of brain tissue — SBEM, ATUM and FIB-SEM — are depicted in Figure 1.

**Figure 1.**
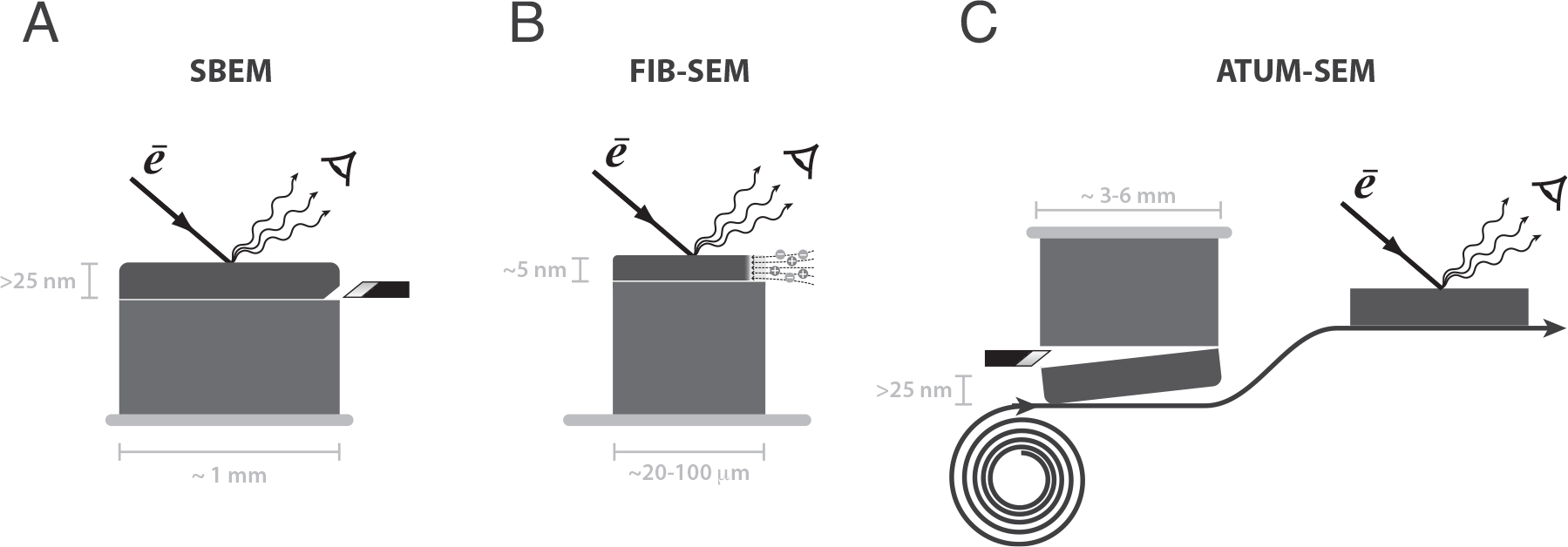
EM connectomics tools: A) Serial block face SEM (SBEM) images the top face of a pre-stained tissue block, then removes the imaged face with a diamond knife, revealing the next layer. B) Focused ion beam SEM (FIB-SEM) operates on a similar principle, but removes tissue layers by ablation with a focused beam of ions. This enables thinner sections and higher electron doses compared to SBEM, but the finite depth of focus of the ion beam limits the size of individual blocks. C) Automated tape collecting ultramicrotomy SEM (ALUM) sections tissue with a diamond knife and places the sections on a solid support, before loading samples into the electron microscope.

### 2.2.1 Serial Block Face SEM (SBEM)

SBEM uses a diamond knife embedded in the SEM to serially remove an ultra-thin section of a pre-stained tissue block [37] after surface imaging, revealing the next layer to be imaged [38].

#### Resolution

The *z*-resolution achievable with diamond knife sectioning is on the order of 25–30 nm, limited by the knife sharpness; note that the section itself can be destroyed in SBEM since it is the block face that is imaged. The effective *z*-resolution of SBEM could be improved by using multi-energy deconvolution SEMs, allowing “virtual sections” thinner than the physical sectioning thickness of the diamond knife [39]^1^. SBEM also imposes a minimal lateral pixel size, since the higher electron doses associated with smaller pixels interfere with reliable physical scraping by the diamond knife when pixel densities surpass this limit [33].

#### Maximum block size

Current implementations of SBEM are limited to tissue blocks ~1 mm on a side, although there appears to be no block size limitation in principle [31].

### 2.2.2 Automated Tape-Collecting Ultra-Microtomy (ATUM)

ATUM [34, 40] allows a block of tissue to be sliced into *>* 25 nm ultra-thin sections which are arrayed on a tape reel for random-access imaging.

#### Resolution

Empirically, the reliability of ATUM-SEM decreases considerably below ∼30 nm section thickness. As for SBEM, virtual sectioning techniques could potentially be used to achieve higher effective *z*-resolution.

Unlike SBEM, ATUM does not suffer from a minimal pixel size limit due to physical tissue damage at high electron doses, since the tissue sectioning occurs *before* imaging. This has allowed a lateral pixel size of 4 nm × 4 nm, such that a voxel size as small^2^ as 4 nm × 4 nm × 25 nm appears to be possible^3^ [41].

#### Maximum block size

ATUM-SEM can achieve large lateral slice sizes, e.g., 2.5 mm × 6 mm, and sufficiently-thin sectioning allows effectively lossless tracing along the axial dimension. Thus, ATUM-SEM appears to be suitable for whole-mouse-brain-scale automation [31].

#### Reliability

Reliability of automated ultra-thin sectioning would likely be the key limiting factor for whole-mouse-brain EM imaging in this approach. One rough estimate gives success rate of 990 per thousand ATUM sections (Richard Schalek, personal communication). In addition, 10000 sections can be cut and collected for each fresh area of the knife (Richard Schalek, personal communication).

### 2.2.3 Focused Ion Beam SEM (FIB-SEM)

In FIB-SEM, a gallium ion beam, rather than a diamond knife, removes a thin layer of the tissue block by ablation [42], to expose a fresh surface for imaging.

#### Resolution

FIB-SEM has achieved 5 nm × 5 nm × 5 nm voxel sizes [42], because it can a) tolerate large electron doses, eliminating the lateral resolution issues of SBEM and b) slice at a very fine *z*-resolution [33]. In fact, the *z*-resolution of FIB-SEM microscopy is limited by depth of electron penetration into tissue block [33], such that lower voltages and more sensitive electron detectors could in principle reduce the slice thickness even further.

#### Maximum block size

The major limitation of FIB-SEM, which appears to be fairly fundamental, is that it can only apply to blocks at most 100 µm across along the direction of the milling beam (with an optimal size of ~20 µm), due to the limited depth of focus within which the ion beam is thin and approximately collimated [33]. Automated FIB-SEM imaging of large volumes of brain tissue would thus involve lossless subdivision of the tissue into rectangular blocks, with one edge length of ∼20 µm and the other edges much longer: for example, blocks of dimensions ∼20 µm × 100 µm × 100 µm might be a reasonable target.

## 2.3 EM Data Acquisition: Cost Estimates

The image-acquisition cost for a 3-year project is given by

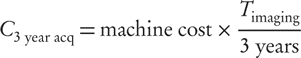

where *T*_imaging_, the time it would take to acquire all the data on a single machine, is given by

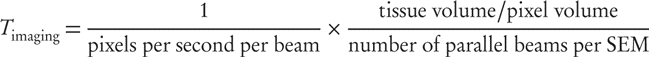

In the below, we typically assume a machine cost of $1M, and compute the imaging time for a 420 mm^3^ brain at the highest achievable resolution on each machine type. Note that if pre-existing machines are used, or if the machine cost can be amortized over a longer duration (e.g., multiple projects), then the effective image-acquisition cost would be lower.

### 2.3.1 SBEM

In one SBEM study, imaging a 325 µm × 325 µm × 60 µm tissue block at 16.5 nm × 16.5 nm × 25 nm voxel size took on the order of 7 weeks at ∼0.5 MHz pixel rate [43]. This is in order-of-magnitude agreement with the simplest calculation, based only on the pixel size and ∼2 µs dwell time: 2 µs × (325 µm × 325 µm × 60 µm)*/*(16.5 nm × 16.5 nm × 25 nm) ≈ 2 µs × 10^12^ pixels ≈ 517 hours ≈ 3 weeks. The estimated cost for a single whole mouse brain acquisition in 3 years is roughly $1B without parallelization and $20M–$100M with 60-fold parallelization. SBEM can likely be operated at lower pixel dwell times (e.g., 0.5 µs) without unacceptable loss of image quality, decreasing the cost proportionately.

### 2.3.2 ATUM

ATUM can achieve 40 megapixel per second imaging rate at 4 nm × 4 nm × 25 nm pixel size (or an effective imaging rate of 400–2400 megapixels per second with 10-to 60-fold parallelization). The estimated 3 year whole mouse brain imaging cost is then $300M and $5M–$30M.

### 2.3.3 FIB-SEM

FIB-SEM can achieve *>* 5 MHz pixel rate at 5 nm × 5 nm × 10 nm voxel size [33]. For a 3-year acquisition, we would need

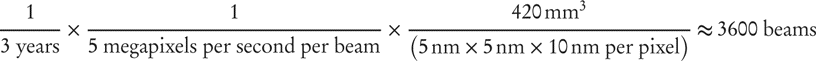

Without parallelization, the estimated 3 year imaging cost is $3.6B, comparable to the estimate of $5B in [33] (which considers more than just imaging costs). At 60-fold parallelization, 60-360 machines would be needed, giving an estimated cost of $60-$360M.

### 2.3.4 Summary

Data *acquisition* costs for whole-mouse-brain automated EM approaches could lie in the range of $10M–$200M. These estimates do not include the costs of *developing* reliable systems for lossless tissue subdivision, thin-sectioning and sample handling.

## 2.4 EM Data Analysis: Basic Properties

A major outstanding challenge in SBEM connectomics is image analysis: reconstructing neuronal wiring from EM image stacks. Tracing thin axons over long distances is the key difficulty, as opposed to synapse detection [30, 44].

### Error propagation

A critical issue is the reliability of the *analysis*. Each error affecting an axon can cause disproportionate damage to the reconstruction, by mis-labeling each of the hundreds of downstream synapses in the connectivity matrix. For example, if an error in an axonal trace occurs on average even once per the length of one axon, which is several mm in mouse brain, then 50% of all connections in the connectivity matrix will be incorrect. In practice, achieving one error per several mm of EM trace is challenging: in one study [45], the errors in the *manual* reconstructions from ssTEM data — i.e., the best reconstruction quality currently available, as compared with automated algorithms — were roughly 1 error per 1000 axonal slices, corresponding to roughly 1 error per ∼50–100 µm of axonal length, far below the ∼4 mm typical axonal length in mouse cortex. In that study, the slice thickness was 50 nm, so decreased error rates would be expected in the techniques studied here, which use *<* 30 nm slice thickness.

### Dependence on voxel size

Currently, the ability of automated algorithms to trace the thinnest axons depends strongly on the imaging resolution. Given appropriate staining, a voxel size of (*<* 10 nm) × (*<* 10 nm) × (*<* 10 nm) is sufficient to allow fully-automated axon tracing, whereas larger voxel sizes can lead to tracing ambiguities that are currently only resolvable through human-assisted image analysis. It is possible, though not proven [31], that a sufficiently small lateral pixel size — e.g., as is achievable in ATUM-SEM due to its tolerance of high electron doses, but not in SBEM — can allow for unambiguous automated neurite tracing even at relatively low *z*-resolutions.

### Dependence on staining method

The quality of EM data depends not only on the instrument resolution but also on the properties of the staining method. Staining of internal structures in axons and dendrites can lead to ambiguities in the resulting images. If only external surfaces are stained (e.g., along with a synapse stain) then even 25 nm × 25 nm × 25 nm instrument resolution may be sufficient for unambiguous axon tracing in some cases. On the other hand, if many internal structures are heavily and non-specifically stained (i.e., the method produces large “blobs” of dense stain), then even 5 nm × 5 nm × 5 nm instrument resolution may not be sufficient for axon tracing. Specific staining of the plasma membrane or other structures using genetically encoded contrast generators (e.g., APEX [46]) may be one option for programmable control of the staining properties. Genetically encoded contrast agents could be targeted to specific neuronal compartments, such as the axon (much as are certain ion channels) [47], in order to sparsify the scene. Reliable and uniform staining of entire mammalian brains prior to tissue sectioning is the subject of ongoing research [37].

### Theoretical limits on the tracing error rate

In EM tracing, the goal is to trace tube-like structures (axons) through a series of images using the fact that the tubes are hollow. The tubes are randomly oriented throughout the series of images, running perpendicular or parallel to the slice with roughly equal probabilities (in cortical neuropil). If the axon is perpendicular to the slice, then it appears as a “circle”. If the axon is oriented parallel to the slice, then it appears as a “blob” of stain arising from its upper and/or lower membrane surfaces. The fundamental parameters are the largest voxel dimension *h* and the smallest opening diameter *d* in the tubes. If two slice-parallel axons nearly overlap, and are heading in nearly the same direction, then their paths cannot be distinguished, even when using longer-range structure across multiple images or sections and even as judged by human experts. This led to a model of the frequency of such “true ambiguities” [45] per micron of axonal wires, as a function of the slice thickness *h*. Using the observed distribution of axon diameters *ρ*(*d*), the model predicts one expected true ambiguity per 100–1000 mm of axonal wire for 20–30 nm sections; recall that there are kilometers of axonal wire per mm^3^ of tissue.

### Data storage requirements

Assuming 10 nm × 10 nm × 10 nm EM voxel size, there are 420 mm^3^*/*(10 nm × 10 nm × 10 nm) ≈5 × 10^16^ voxels in a 420 mm^3^ mouse brain. At 1 byte per pixel, this is ∼400 000 terabytes of EM image data, roughly the total amount of data transmitted over the internet during a 10 hour period circa 2013 (storage would cost $20M on $100 2TB hard drives).

## 2.5 EM Data Analysis: Cost Estimates

### 2.5.1 SBEM and ATUM

The standard *z*-resolution of SBEM and ATUM of 25–30 nm is not sufficient to allow fully-automated tracing of neuronal processes with currently available algorithms. Manual volume segmentation from SBEM image stacks by a trained human requires roughly 2 work-hours per µm^3^. To get around this, Helmstaedter and colleagues [30] split the analysis pipeline into two separate stages: skeleton tracing and volume segmentation / contact detection.

For the skeleton tracing step, REdundant-Skeleton COnsensus Procedure (RESCOP) [48] is a human-assisted process for tracing the center of the axon. The software resolves disputes between users through redundancy and infers an estimate of the skeleton trace via a statistical model. A redundancy factor of 18 or 19 leads to roughly one tracing error per cell. This method achieved ~0.0135 work-hours per µm^3^. At a labor rate of $5 per hour, this corresponds to $70M per mm^3^; for the whole mouse brain the labor cost would be of order $30B. To complete the analysis within 3 years using this method, assuming 2000 working hours per year, 945 000 laborers would be required.

The human-assisted skeleton tracing does not reveal synapses or detailed local morphology. This information is obtained via fully-automated volume segmentation algorithms, applied after the skeleton tracing [49, 50]. The estimated volume error rate for this process is around 3% [30]. Note that this procedure currently does not reveal “ground truth” synapses as defined by the presence of a post-synaptic density (PSD) and pre-synaptic vesicles, but merely assesses the probability of connected neurons based on the pattern of contact between two cells (e.g., contact area, which is not a good predictor of actual synapses [30, 51], except at very high contact areas [30]).

In an alternate workflow, segmentation can be performed automatically, followed by human proofreading [11]. Assuming (15 nm)^3^ voxels, a recent review [11] estimated that current methods would require 4.5 million person years of proofreading for a whole mouse brain, similar to the 3 · 945000 = 2.84 million person years estimated above for manual skeleton tracing. Thus, either segmentation algorithms must be improved, or data quality must be improved to compensate, to allow a dramatic reduction in the need for either pre-segmentation manual skeleton tracing and/or post-segmentation manual proofreading.

Large-scale internet-based crowd-sourcing could play an important role in scaling up data analysis, since tens of thousands of users appear to be willing to participate in the process for “free” [52]. These players also collectively generate a large data-set for training machine learning algorithms [52]. Other crowd-sourcing approaches for image segmentation are also being developed [53].

Using today’s tools, analysis costs would be in the tens billions of dollars for a whole mouse brain. The computational connectomics sub-field aims to reduce the analysis costs by orders of magnitude, ideally leading to full automation, and it is making progress towards this goal [54, 55].

### 2.5.2 FIB-SEM

It is possible that the (*<* 10 nm) × (*<* 10 nm) × (*<* 10 nm) resolution of FIB-SEM will enable reliable, fully automated axon tracing and synapse identification from large volumes [33]. Automated synapse detection from FIB-SEM images has been demonstrated with error rates comparable to that of human experts (e.g., 0.92 recall at 0.89 precision) [56, 57].

## 2.6 Annotation of EM Connectomes

While stains have been developed to couple electron-imaging contrast to neuronal and vesicular membranes, there are few extant mechanisms to couple electron contrast to other forms of sub-cellular molecular information, such as specific genetic sequences or specific proteins. Recent attempts have been made to introduce multiplexed labeling capabilities into EM [58], as well as to create genetically encoded proteins which can serve as EM markers [46, 59]. Furthermore, it may be possible to create nanoscale spatial patterns of heavy metals or other high-contrast elements which could serve as combinatorially-diverse EM labels (EM barcodes). Another option for obtaining multiplexed molecular information from a given cell body would be as follows: given the > 1000 sections that contain a single cell body, it would be possible to antibody-stain each section for a different molecular marker, and thus to assign a “molecular identity” to every EM-reconstructed cell, without requiring any single EM image to be “multi-colored”. Nevertheless, EM currently lags behind optical microscopy in the ability to readily reveal biochemical information in a multiplexed fashion and in any neuronal compartment.

## 2.7 Summary

Electron microscopy imaging using serial block-face SEM (SBEM), automated tape-collection lathe ultramicrotomy (ATUM) or focused ion beam SEM (FIB-SEM) would cost hundreds of millions to billions of dollars for whole-mouse brain data acquisition using current instruments. Next-generation parallel-beam SEMs — e.g., a 61-fold parallelized SEM under development by Zeiss — could reduce the data-acquisition costs into the range of tens of millions of dollars or below, depending on the degree and cost of parallelization.

FIB-SEM will likely allow fully-automated image analysis, due to its *<* 10 nm *z*-resolution and compatibility with 5 nm in-plane resolution. However, due to its limited field of view per instrument (∼20 µm along the milling axis), new instrumentation would be required to automate the sub-division of tissue into appropriate-sized blocks. Hayworth has demonstrated preliminary proof of principle that this sub-division could be achieved without information loss, to enable tracing of fine axons between blocks. SBEM and ATUM-SEM are more readily automated on the hardware side than FIB-SEM due to their compatibility with larger fields of view.

For SBEM and ATUM, which have *z*-sectioning limits of ∼25 nm, tracing of fine axons becomes more difficult for current image-segmentation software. Recent software advances, which separate skeleton-tracing (human-assisted) from subsequent volume segmentation and synapse identification (automated), have reduced the human labor requirements to roughly one work-minute per cubic micron (although current semi-automated image analysis methods mandate a staining protocol incompatible with “ground-truth” synapse identification, i.e., the presence of vesicles and PSD). At a labor rate of $5 per hour, analysis of a whole mouse brain using this software would cost tens of billions of dollars and require nearly a million workers. Further advances in software are needed, therefore, to enable fully-automated analysis of image data generated from SBEM and ATUM. Importantly, the analysis costs could ultimately become negligible, in principle, through algorithmic advances. Also, the effective *z*-resolution of SBEM or ATUM could be improved through virtual sectioning.

Thus, given either a) construction of an automated tissue sub-division system for FIB-SEM or b) full software automation of SBEM or ATUM image analysis (e.g., via machine learning advances), *and* the emergence of multi-beam SEMs at a cost comparable to current single-beam SEMs, a whole mouse brain EM connectome project could be achievable for a cost of tens to hundreds of millions of dollars and a duration of several years per mouse brain. A major advantage of EM connectomics is its ability to trace in detail the morphology and compartmental structure of neurons, which is tightly coupled to their electrochemical functions [60].

## 3 Trans-Synaptic Barcode Pairing and Bulk Sequencing (BOINC)

A DNA barcode is a unique sequence of DNA used to “tag” an object of interest. Zador has suggested [3] an approach to connectomics, called Barcoding of Individual Neuronal Connections (BOINC), which leverages large numbers of DNA barcodes. First, each neuron is given a unique DNA barcode. Copies of each neuron’s barcode are then exchanged with its immediate synaptic neighbors. A cell’s own barcodes are then stitched together with barcodes received from its synaptic neighbors, forming a set of barcode pairs corresponding to synaptically connected neurons. Zador’s original proposal suggested one potential implementation: using trans-synaptic tracer viruses (e.g., engineered pseudorabies replicons) to shuttle copies of the barcode from a given cell to its immediate presynaptic neighbors, whereupon a recombinase (e.g., phiC31 integrase) in the recipient cell would link donor and recipient barcodes into a single strand [3].

The barcode-pair DNA strings from all cells are extracted, pooled, amplified (i.e., creating many copies of each barcode pair) and sequenced on a bulk DNA sequencing machine, such as an Illumina HiSeq. This results in digital data specifying a set of “on” matrix elements, corresponding to barcode pairs (synaptic neighbors) which are observed, and a set of “off” matrix elements, corresponding to barcode pairs which are not observed (e.g., due to the absence of a synapse between the corresponding two neurons).

To allow “annotation” of the connectivity matrix, Zador and colleagues also suggested that additional information, encoded in nucleic acids, could be appended onto these barcode pairs, e.g., RNA sequences indicative of a cell’s gene expression profile (cell type).

Note that the problem of determining the *spatial position* of each neuron is not solved by this approach, although coarse-grained positional information could be included by sectioning the tissue and appending additional, position-encoding DNA barcodes to the cell-barcode pairs extracted from each physical section, prior to bulk sequencing. The basic idea of BOINC is depicted in Figure 2.

**Figure 2.**
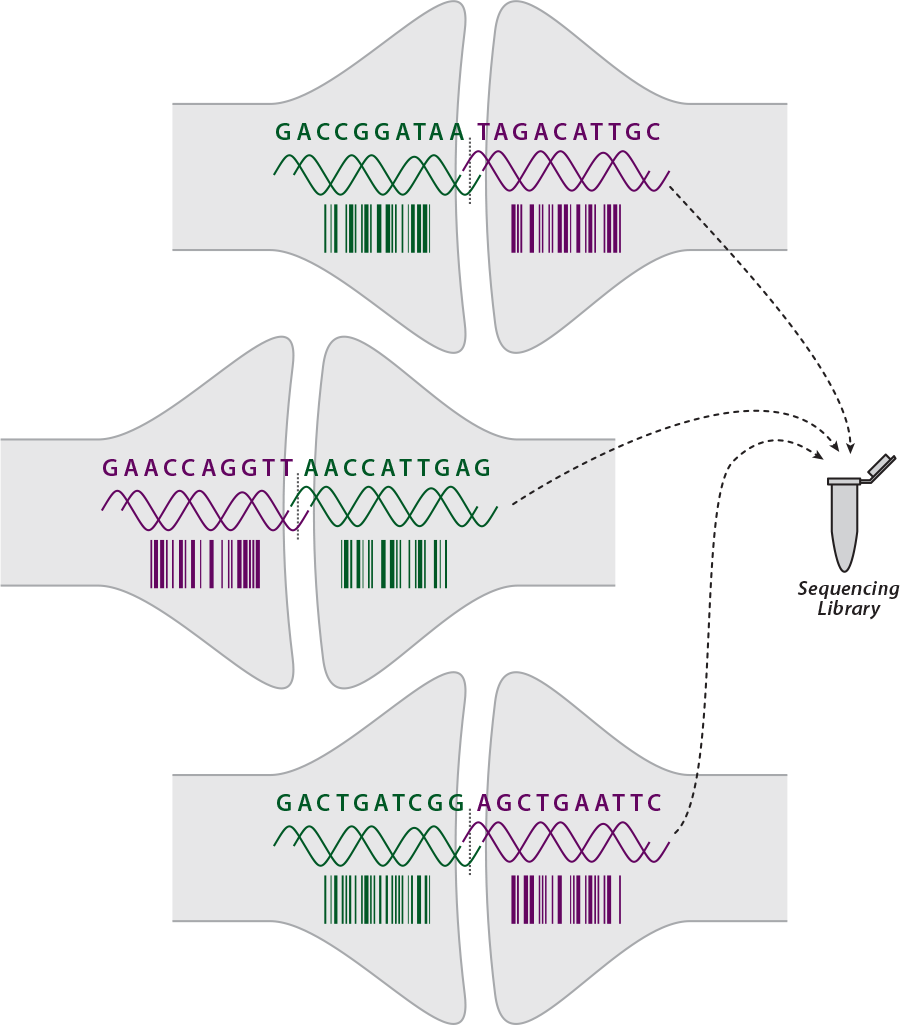
Reading out neuronal connectivity via bulk sequencing: cell-identifying nucleic acid barcodes from synaptically-neighboring cells are physically linked (e.g., via viral exchange and recombinase activity [3] or other methods [63]), and extracted from the neural tissue. The linked barcodes are then sequenced on a high-throughput DNA sequencer, such that each sequencing read corresponds to a barcode pair from a synaptically-connected pair of neurons.

Alternate molecular implementations of the same idea (e.g., which obviate the use of trans-synaptic viruses [61]) could be preferable from a practical standpoint. For example, synaptoneurosomes containing cell-specific barcode RNAs could be extracted from the tissue and their contents sequenced via a vesicle-barcoded emulsion PCR: synaptoneurosomes typically have some of the pre-synaptic and some of the post-synaptic membrane still attached and even re-sealed [62], although there would be an issue of synaptoneurosome collection efficiency in this scheme.

### 3.1 DNA Barcodes

In one implementation, the DNA barcodes are contiguous strings of random nucleotides (random oligonucleotides) [64, 65]. In another implementation, the barcodes correspond to an array of direct or inverted DNA sub-strings flanked by recombinase inversion sites [3] (e.g., with 19 nucleotide inversion sites for Rci recombinase [66]). The stochastic arrays could be generated in-vivo by recombinase activity, starting from a standard cassette present in all neurons. There is precedent for recombinase-based sequence diversity generation in biology: the Min system makes 240 distinct variants of its multiple-inversion site, leading to 240 different isomeric forms of a phage coat protein to evade bacterial defenses [67].

In the first implementation, DNA barcodes consisting of only 20 DNA nucleotides (A, T, C or G) could in principle uniquely label 4^20^ = 10^12^ neurons, four orders of magnitude larger than the number of neurons in a mouse brain. When barcodes are generated (or chosen) randomly, there is a need to consider the probability of two neurons acquiring the same barcode. To uniquely identify a cell with a DNA barcode, the barcodes must be long enough to avoid the occurrence of duplicate barcodes in the population. The probability of no identical barcodes when *n* barcodes are chosen with replacement from a test-tube with 4*^j^*barcodes (i.e., with all possible DNA oligonucleotides of length j) is

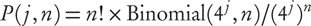

where *n* is the size of the cell population and *j* is the DNA barcode length in nucleotides [68].

For *n* = 7.5 × 10^7^ neurons and *j* = 31 base-long barcodes, the probability of a duplication 1 - *P* (*j*, *n*) *<* 0.001 (the per-neuron probability of duplication is then roughly 10^−11^). This corresponds to a total barcode population size of 4^31^ ≈ 5 × 10^18^.

For the case of recombinase inversion barcodes, the number of barcodes generated from *k* segments is *k*! × 2^*k*^, as long as the recombinase inverts but only rarely excises on the relevant timescales [3, 69]. To achieve a similar probability of barcode duplication, only *k* ≈ 16 distinguishable segments are needed.

There are many other strategies to create cell-identifying barcodes besides the two just mentioned; the diverse mechanisms involved in generation of antibody diversity by the immune system provide a range of examples. Indeed, somatic (VDJ) recombination has been used as a form of in-vivo barcoding for tracing of lymphocyte lineages in the mouse [70].

**Error sources** PCR amplification and sequencing can introduce errors which would transmute one barcode into another. Fortunately, the recombinase-based barcode generation strategy leads to barcodes that are highly orthogonal at the sequence level (large minimal pairwise edit distance between barcodes, compared to the mutation probability), and synthetic barcode libraries introduced via viral transduction could be designed to be highly orthogonal. On the other hand, short barcodes strings which are generated stochastically in all cells by other methods will not necessarily be highly orthogonal.

Illumina paired-end sequencing can achieve error rates of roughly *p* = 0.01^2^ = 10^−4^ per base. Assuming a 100 bp template, the probability of two errors is then *p*^2^ · Binomial(100, 2) = 5 · 10^−5^. The error rate per cycle of PCR is much lower due to the high fidelities of proofreading polymerases: *f* = 5 · 10^−7^ per base for Pfusion [71]. The fraction of strands with ≥ 1 polymerase-induced error after *d* cycles of PCR on a template of length *b* nucleotides is then *F* (≥ 1) = 1 - *e*^−*b*·^ *^f^* ^·*d*^ = 0.00125 [72] for *d* = 25 cycles and *b* = 100 nucleotides. On the other hand, in complex template libraries, errors due to mis-priming and chimeric products can occur at rates of 5% or higher. It is possible to reduce the effective PCR and sequencer error rates using “digital” sequencing methods like [71, 73], which employ pre-amplification template barcoding and redundant sequencing to factor out these error sources.

Failure to capture any barcode pair corresponding to a given connection, leading to a false negative (missed connection) in the connectivity matrix, will likely be the dominant source of error in most implementations of BOINC. With highly orthogonal barcode sequences, false-positives due to sequencing errors can be minimized. Therefore, it is likely possible to implement BOINC in a regime where almost all errors are false-negatives, in contrast to the electron microscopic axon tracing approaches which are quite vulnerable to false-positives [3].

### 3.2 High-Throughput DNA Sequencing

The cost for a BOINC connectome is *C*_BOINC_ = *c* · *r* · *N*_synapses_ where *c* is the cost per sequencing read, *r* is the number of sequencing reads per synapse and *N*_synapses_ is the number of synapses in the tissue under study. The fraction of un-sampled synapses is *f*_unsampled_ = *e*^−^*^r^*[3] so that 1 - *e*^−10^ = 99.995% of synapses are sampled at *r* = 10 and 95% of synapses are sampled at *r* = 3. Because many pairs of neurons are connected by several synapses, the fraction of un-sampled connections (synaptically linked cell pairs) will be less than the fraction of un-sampled synapses.

The mouse brain contains roughly *N*_synapses_ = 10^11^ synapses: an average of 10^3^ synapses per neuron gives *N*_synapses_ = 7.5×10^10^, whereas an approximate average spatial density of 1 synapse per µm^3^ gives *N*_synapses_ = 4.2×10^11^. Hence 10^11^–10^12^ sequencing reads are required per mouse connectome, depending on the redundancy factor *r*.

With current sequencing technology, running 3 lanes of an Illumina HiSeq 2500 produces *>* 10^9^ reads (of up to 100 bp each) in about 10 days for a cost of a few thousand dollars. Roughly 100 HiSeq runs would be required for a full mouse connectome, for a cost of a few hundred thousand dollars. An existing high-throughput genomics facility (with *>* 50 HiSeq machines) could sequence a mouse connectome in 1-2 months.

The cost per base-pair (bp) of DNA sequencing has been decreasing rapidly: 2 bp per dollar in 2004, 10^6^ bp per dollar in 2009 and 10^7^ bp per dollar in 2011 [3, 74]. The “$1000 human genome” corresponds to $1000/(3 · 10^9^ bp · 40×) = $10^−8^ per bp, assuming 40× coverage. At these rates, the cost per 100 bp read is $10^−6^. Thus the minimum cost at these rates is about $10^−6^/synapse, or about $100k for 10^11^ synapses. Three-fold and ten-fold oversampling (*r* = 3 or *r* = 10) raise the cost to $300k and $1M per whole mouse brain, respectively. Corresponding costs for the mouse cortex alone, which contains perhaps 10% of all synapses, range from $10k to $100k.

If these trends continue, it is not unreasonable to imagine that sequencing costs for a mouse brain connectome could drop by a further factor of 10 or more in the foreseeable future. At that point other expenses, including mouse and DNA processing costs, will dominate. Note that we have not included the cost of the bulk sequencing machines in this calculation: we are assuming that existing machines are used, e.g., at an existing genomics facility.

### 3.3 Annotation of BOINC connectomes

At 100–200 bp, each sequencing read would have enough room to include a minimal amount of transcriptomic information, in addition to just the connectivity matrix. This could take the form of RNA transcripts attached to the barcodes via RNA transsplicing. Quantitating the relative proportions of just a few transcripts could be useful: for example, GAD67 and NeuN can be used to identify inhibitory neurons [75]. Sequencing and abundance-counting of a few dozen transcripts could be sufficient to identify known neurobiologically relevant cell types: PV, SOM and VIP to identify the major classes of interneurons, for instance, and DAT, CHAT and others to identify major classes of neurotransmitter-secreting cells. Reliably implementing such trans-splicing mechanisms may be difficult in practice, however, and the method does not scale to capture full transcriptomes. As an alternative, BOINC connectomes could be annotated with transcriptional information via cell-specific barcoding of ribosomes [61].

It is also possible that relative connection strength annotations could be incorporated into BOINC by counting the number of recovered barcode pairs corresponding to any given pair of cells. In many potential implementations of BOINC, the number of barcode pairs recovered from a given cell pair would scale approximately linearly with the total area of synaptic contact between the cells, which may be correlated with connection strength [76–78], although the precise extent to which this relation holds is not known and some potentially complicating factors have been identified [79]. Variability in the barcode pair collection efficiency across different cells could confound such measurements, however, and total contact area is likely not a perfect indicator of connection strength.

While BOINC can also be annotated with coarse-grained positional information, its major limitation is that it does not reveal the precise spatial position or morphology of each cell. Optical microscopy techniques incorporating BOINC barcodes could potentially ameliorate this, as discussed below.

## 4 Direct Optical Microscopy for Connectomics

An optical microscopy approach to connectomics would be powerful, in principle, in that it could allow integration with a wide range of other biochemical measurements that are accessible through modern light microscopy, e.g, Fluorescent In-Situ Hybridization (FISH) [80, 81] or serial histology [82, 83]. It is widely believed, however, that electron microscopy is the only approach which can allow acquisition of connectomes by direct imaging. Indeed, there can be as many 10-40 neurites per diffraction-limited optical resolution volume [12], which creates severe difficulties with direct optical tracing of axons, even when neurites are tagged with distinct sets of fluorescent proteins through random genetic recombination (BrainBow) [84–86]. Nevertheless, there may be novel strategies which can work around this limitation.

### 4.1 Observing Synapses vs. Tracing Axons

Because of the comparative sparseness — at 1-2 synapses per µm^3^ — of synapses in 3D space, optical connectomics approaches could succeed by restricting their attention *only* to the synapses themselves [12]. Rather than directly tracing the paths of axons and dendrites through a series of images, cell-identifying molecules could be physically trafficked — via endogenous cellular processes — to the pre-synaptic and post-synaptic compartments [87–89]. Then, *observations of the synapses alone* could reveal the identities and/or properties of the pre-synaptic and post-synaptic cells.

#### Resolution requirement to resolve neighboring synapses

Diffraction-limited 3D imaging (*λ/*2NA ≈ 200 nm *xy*-resolution and 2*λ/*NA^2^ ≈ 533 nm *z*-resolution for numerical aperture NA = 1.5 and wavelength *λ* = 600 nm) is not sufficient to directly resolve a synapse from its neighboring synapses [12]. Simulations of synapse-labeled fluorescence microscopy based on EM reconstructed rat hippocampal neuropil have suggested, however, that *<* 100 nm isotropic resolution is sufficient to resolve >90% of synapses from their nearest neighbors [12]. These simulations assumed that fluorescence was limited to the pre-synaptic and post-synaptic densities (PSDs), as opposed to the entire axonal bouton or spine head.

Figure 3 shows a conservative estimate of the resolvability of nearest-neighbor synapses based on the dataset from [12], in which synapses are present at an average density of 1.85 per µm^3^. A strict criterion for resolvability is applied: two synapses are considered to be non-resolved if any of their labeled points are separated by a distance smaller than the isotropic resolution. Since synapses are extended objects, it is often possible to separate them based on shape, even if they are not resolvable according to the strict criterion; the strict criterion gives a sufficient but not necessary condition for resolvability.

**Figure 3.**
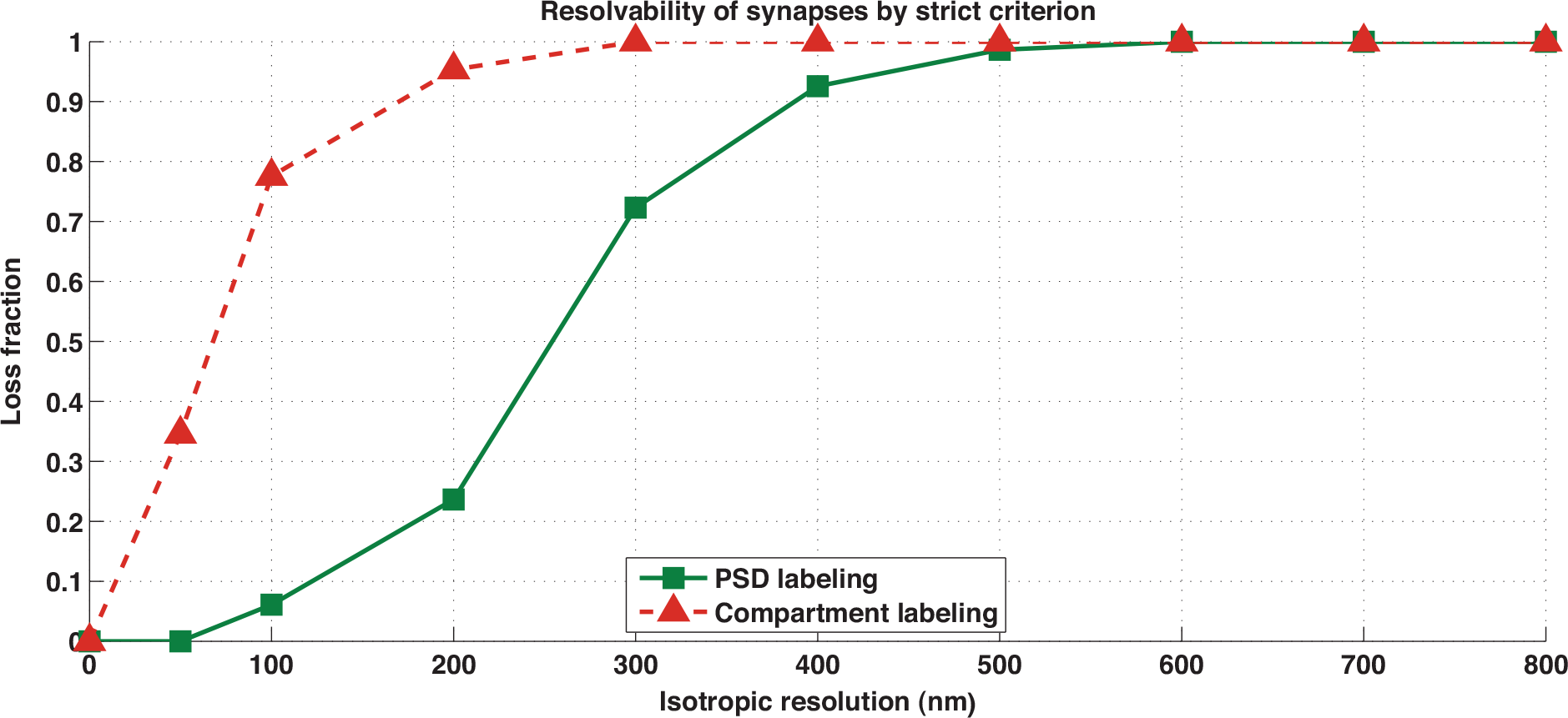
Optical resolution requirements for resolving nearest-neighbor synapses. The fraction of non-resolved synapses as a function of isotropic resolution for PSD labeling (green) and whole-compartment labeling (red), based on the dataset from [12]. A pair of synapses is considered unresolved here if and only if they contain labeled points separated by less than the isotropic resolution.

Labeling only of the PSDs allows resolution of >90% of synapses at isotropic resolution *<* 125 nm, whereas labeling of the entire pre-synaptic and post-synaptic compartments gave poor performance even at *<* 50 nm isotropic resolution. The poor performance for whole-compartment labeling is not surprising: synaptic boutons and spine heads often directly contact other nearby boutons and spine heads, leading to high confusion rates between nearby synaptic puncta, in the whole-compartment labeling scenario, even if the imaging resolution were to approach to zero. Therefore, to optically resolve individual synapses, it is essential that the labeling be highly specific to the PSDs, as could perhaps be achieved with a protein-tagging strategy.

#### Achieving the required resolution

Experimentally, confocal microscopy in *<* 100 nm thin sections and at roughly 200 nm diffraction-limited *x y* resolution — in the context of Array Tomography — appears to optically resolve most if not all synapses [82, 83, 90] via antibody staining of synaptic proteins such as synapsin. Isolated fluorescent puncta are observed, in numbers similar to those expected in the tissue based on EM measurements of synapse density [83]. In one recent study, the fluorescent puncta have been attributed to individual synapses [91] by comparison with EM imaging of the same serial sections.

Advances in microscopy could minimize the need for ultra-thin 2D sections. The dual-objective imaging technique *I* ^5^*M* achieves 100 nm resolution axially and 200 nm resolution laterally in a wide-field mode [92], and multi-photon 4Pi-confocal microscopy gives similar axial resolution [93] in a parallelized beam-scanning mode.

A 10–100× improvement to the speed of linear structured illumination microscopy (SIM) has recently been reported [94]. Linear SIM exceeds the diffraction-limited resolution by a factor of 2 along all three axes, with commercial systems achieving 130 nm × 130 nm × 270 nm resolution voxels. Further improvement to the *axial* resolution of SIM could allow it to resolve most synapses. For example, *I* ^5^*S* two-objective detection [95] is a form of SIM with isotropic 100 nm resolution.

Other techniques offer even deeper levels of optical super-resolution. Nonlinear SIM–SIM performed at illumination intensities high enough to saturate the fluorophore – can improve resolution beyond that of linear SIM [96], and parallelized nanoscopies based on point-spread function engineering have been demonstrated [97]. Stochastic Optical Reconstruction Microscopy (STORM) achieves 30 nm × 30 nm × 50 nm voxel size in 3D [98], but at its current volume throughput of roughly 15 µm^3^/s, STORM of an entire mouse brain would take nearly 1000 imaging years.

Molecular methods could be used to increase the effective spatial resolution, relative to that of any given optical setup, by “stratifying” the observation of different synapses into different imaging frames [99]. This would increase imaging time proportionately. For a 2× cost in the imaging time, molecular stratification could also resolve the pre-synaptic and post-synaptic compartments of a given synapse: first activate pre-synaptic but not post-synaptic dyes, then switch to a new camera frame and reverse the activation pattern.

## 4.2 Strategies for Optical Connectomics

### Fluorescent protein-based synaptic BrainBow

A “synaptic BrainBow” strategy [12] has been proposed, in which each cell would express a distinct *combination* of fluorescent proteins, which would be targeted to the pre-synaptic and post-synaptic compartments. Then, by observing the spectrum of colors at each synapse, the corresponding pre-synaptic and post-synaptic cells could be identified, even if the pre-synaptic and post-synaptic compartments of a given synapse are not optically resolvable from one another. This could be combined with observation of the corresponding fluorescent protein color patterns expressed in the nuclei, thus labeling the locations of the corresponding somas.

This method could have favorable properties with respect to resolution of neighboring synapses, outperforming the conservative resolution requirements in Figure 3. In particular, synaptic BrainBow relies on tagging synapses based on co-localization (spatial correlation) of fluorescence from pre-synaptic and post-synaptic markers: even if the fluorophores are not precisely localized to the pre-synaptic and post-synaptic densities, their emissions co-localize only over the synaptic cleft itself. Therefore, detection based on fluorescence co-localization can perform better than directly resolving single-colored synaptic puncta.

Unfortunately, the originally-proposed form of synaptic BrainBow [12] does not scale to entire mouse brains because of the limited color palette of available fluorescent proteins: 2 · log_2_(10^8^) = 54 spectrally distinguishable fluorophores would be required [12].

### Fluorescent In-Situ Sequencing (FISSEQ) for 4*^N^* -“color” synaptic labeling

Novel methods could potentially allow variants of the synaptic BrainBow strategy to scale to mammalian systems. An alternative method could leverage Fluorescent In-Situ Sequencing (FISSEQ) [63, 99], a recently-developed method for sequencing of DNA or RNA by optical microscopy in the context of intact tissue slices. In effect, FISSEQ constitutes a form of fluorescent microscopy in which there are 4*^N^* distinguishable labels, corresponding to the 4*^N^* possible nucleotide sequences of a DNA molecule of length *N* nucleotides. By leveraging FISSEQ, it may therefore be possible to create a 4*^N^* -“color” variant of the synaptic BrainBow strategy, which would scale readily to whole mouse brains, despite using only four *actual* spectrally distinguishable fluorophores. In one possible implementation, cell-identifying RNA barcodes (similar to those used in BOINC) could be targeted to the pre-synaptic and post-synaptic densities, and their nucleotide sequences could be read out by fluorescent microscopy in-situ.

If the fluorescent sequencing frame rate of an Illumina HiSeq machine^4^ were directly translated to in situ sequencing of 100 nm thick tissue slices in a diffraction-limited microscope, similar to the setup used in Array Tomography [82, 83], the imaging time^5^ and imaging cost for a 3-year mouse brain connectome would be

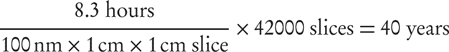

and $13M respectively, assuming $1M per Illumina-rate machine.

## 5 Technology Development Pathways

These approaches could be validated in smaller brains. For example, the *Drosophila* brain, with 135k neurons, is roughly 1000× smaller than the mouse brain. In the electron microscopy approaches, only a few microscopes would be required for *Drosophila*, although image analysis would still pose significant challenges.

For BOINC, a single 11 day run on a HiSeq produces *>* 10^9^ reads, more than sufficient for a *Drosophila* connectome (e.g., 10^8^ synapses × *r* = 10 reads per synapse). Reads of length 100 bp could include two 20-base barcodes, to uniquely label all neurons in the fly, as well as additional barcodes to provide spatial information. Indexing 10 sections along the *x*, *y* and *z* axes – forming blocks of *<* 100 µm edge length – would require only log_4_(10^3^) = 5 additional nucleotides, or *<* 10 additional nucleotides for a highly orthogonal set.

For an optical microscopy approach based on in-situ sequencing of synapse-localized RNA barcodes, roughly 5000 *z*-sections of 75 nm thickness and 400 µm × 1000 µm *xy* cross-section would be sufficient to cover the entire *Drosophila* brain. The totality of these sections would fit on a single standard microscope slide. If a 4-color 2D saturated SIM [96] image at 50 nm *xy* resolution takes 1 s to acquire and comprises a 50 µm × 50 µm field of view, then the time to image all the slices from a single fly is roughly 9 days. This is multiplied by a factor of 20 to account for 20 FISSEQ cycles. Therefore, ultra-thin-sectioning 2D SIM FISSEQ of an entire Drosophila brain at 50–100 nm × 50–100 nm × 75 nm resolution – likely sufficient to resolve nearly all synapses – could be performed in *<* 6 months on a single automated SIM microscope.

Once validated in a smaller model organism, extension to mammalian systems could be straightforward, although different model systems pose different obstacles for genetic engineering tasks like whole brain cellular-resolution barcoding. In addition, technologies like bulk EM staining may need to be adapted [37] to larger volumes. Due to its small brain size, with only a few million cortical neurons [106], the Etruscan shrew may be a desirable early target.

## 6 Summary

Several approaches for whole-mouse-brain connectomics may be nearly within reach for roughly $100M–$200M in a three-year project. For electron microscopy approaches, this would require dramatic improvements in the speed and accuracy of computerized axon tracing. Improvements to the reliability and automation of electron microscopy sample handling would also be essential.

Approaches leveraging a new “exponential resource” — nucleic acid sequence-space — appear to have the potential to further reduce the cost by a factor of 10–100 or more. For example, BOINC [3], a set of approaches based on bulk sequencing of nucleic acid barcodes that have been exchanged across the synaptic cleft and physically paired into a single sequencing read, could potentially obtain a mouse connectome for under $1M at today’s sequencing costs. Further cost reductions are anticipated given the exponential improvement of DNA sequencing technology [74].

More speculatively, the ability to measure combinatorially-multiplexed molecular information (the 4*^N^* possible RNA sequences of length *N*) in situ via optical microscopy, and to localize this readout specifically to synapses, could enable optical microscopy to directly acquire connectomes from fixed tissue samples. This approach could be feasible in the $10M range via a suitable combination of fast super-resolution microscopy [94, 96, 97], physical and/or optical thin-sectioning microscopy [82, 83, 95, 104, 107] and molecular stratification techniques.

The development of a whole mammalian brain connectomics capability will be a significant engineering challenge, regardless of the technology platform(s) adopted. Even once the component technologies are developed, there will be a need to integrate components into an automated pipeline for connectome acquisition. This is most likely to take place if technological innovations enabling significant cost reductions are introduced as early as possible.

## 7 Acknowledgments

Josh Glaser and Ben Stranges for discussions on barcodes, Todd Huffman for discussions on serial sectioning, and Ken Hayworth and Richard Schalek for discussions on EM automation. Dario Amodei, Juan Batiz-Benet, Ted Cybulski, Tom Dean, Noah Donoghue, Kevin Esvelt, Russell Hanson and Jason Pipkin for discussions.

Adam Marblestone is supported by the Fannie and John Hertz Foundation fellowship. Ed Boyden is supported by the National Institutes of Health (NIH), the National Science Foundation (NSF), the MIT McGovern Institute and Media Lab, the New York Stem Cell Foundation Robertson Investigator Award, the Human Frontiers Science Program, and the Paul Allen Distinguished Investigator award. David Dalrymple is supported by the Thiel Foundation. Reza Kalhor, Evan Daugharthy, Seth Shipman and George Church acknowledge support from the Office of Naval Research and the NIH Centers of Excellence in Genomic Science. Evan Daugharthy is also supported by the NSF Graduate Research Fellowship (DGE1144152) and Seth Shipman by the National Institute on Aging (5T32AG000222-22). Konrad Kording is funded in part by the Chicago Biomedical Consortium with support from the Searle Funds at The Chicago Community Trust. Yuriy Mishchenko acknowledges support from Bilim Akademisi – the Science Academy, Turkey, under the BAGEP program, and from BAP Scientific Research Projects Fund of Toros University. Tony Zador, Ian Peikon and Justus Kebschull acknowledge an NIH TR01 award and the Paul Allen Distinguished Investigator award. Justus Kebschull acknowledges a PhD fellowship from the Boehringer Ingelheim Fonds.

## Footnotes

1 Virtual sectioning has particular application to reset sections (the first sections acquired after resetting the cutting arm of the ultra-microtome. ThruSight (FEI, Co) is a commercial application of this idea.

2 The ATUM approach has been routinely applied to image at pixel resolutions down to 1 nm for the imaging of c-elegan neural processes; even sub-nanometer pixel resolutions are possible, but this is slow and in most cases can be considered as oversampling (Richard Schalek, personal communication).

3 Connectivity and synapses may be visible even with an 8–10 nm pixel and proper staining (Richard Schalek, personal communication). In this case, imaging time and imaging cost decrease by a factor of 4. Furthermore, ATUM-based imaging has demonstrated the ability to perform multi-scale resolution (e.g., large axon tracts can be imaged at one pixel size and dwell time, while neuropil can be imaged at a smaller pixel size and dwell time); this further decreases the imaging time and cost.

4 Illumina machines can achieve cluster densities on the sequencing flow cell (essentially a glass microscope slide) of 1,000,000 clusters per mm^2^, similar to the *areal* density of synapses in a 0.5–1 µm thick tissue section. Given that a HiSeq run takes roughly 250 hours (11 days) and generates 300 billion bases of sequence (e.g., 3 billion 100 bp reads), the time to sequence a 1 cm^2^ area is 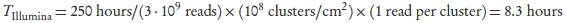

5 For comparison, whole mouse brain fluorescence Micro-Optical Sectioning Tomography (fMOST) at 0.6 × µm 0.8 × µm 1 µm *xyz* voxel size took 19 days [100–105]. This is broadly consistent with the estimate given here for whole mouse brain FISSEQ at the Illumina scan rate and Array Tomography slice thickness: using 100 nm rather than 1 µm sections gives a factor of 10 relative to fMOST, and the 30 cycles of FISSEQ give an additional factor of 30, leading to 15 imaging years for whole mouse brain at the effective throughput of fMOST.

